# IRES-TrAPPr reveals novel insights into viral and cellular mRNA translation

**DOI:** 10.64898/2026.05.06.723280

**Authors:** Gemma E. May, C. Joel McManus

## Abstract

Ribosome recruitment to human mRNAs is thought to occur primarily via cap-dependent initiation (CDI). This process is suppressed during a variety of cellular stresses, including viral infection, suggesting stress-response genes and viral mRNAs use alternative mechanisms to initiate translation. Indeed, many viruses recruit ribosomes directly via Internal Ribosome Entry Sites (IRESes). Hundreds of human mRNAs have been reported to also contain IRESes due to their ability to enhance expression in bicistronic and backspliced circRNA plasmid reporters. These DNA-based screens also reported hundreds of novel IRESes from more than fifty human viruses. However, such assays are prone to false-positives due to promoter and splicing activity, do not compare CDI and IRES translation, and lack the temporal resolution necessary for stress-response studies. To address these issues, we developed IRES-<u>Tr</u>anslating <u>A</u>ffinity <u>P</u>rotein <u>Pr</u>ofiling (IRES-TrAPPr), a massively parallel reporter assay that simultaneously quantifies CDI and IRES activity from thousands of co-transfected mRNAs. We validated this new method using luciferase assays and structure-function analyses of established viral IRESes, demonstrating exquisite sensitivity and specificity. Using IRES-TrAPPr, we quantified the activities of IRES elements from hundreds of viruses from a diversity of hosts. Our results provide evidence that viral IRESes from warm-blooded hosts have adapted higher structural stability to maintain folding at higher temperatures. Finally, we find hundreds of candidate human and viral IRESes from DNA-based screens have negligible IRES activity. Altogether, our results show that IRES-TrAPPr provides a novel, accurate platform for IRES research.

## Introduction

Protein synthesis via mRNA translation is essential for life. mRNA translation can be separated into three processes - initiation, elongation, and termination. In eukaryotes, translation initiation is thought to occur primarily through recruitment of a pre-initation complex (PIC) to mRNA 5’ ends, followed by a directional (5’ to 3’) search for appropriate AUG start codons^1,2^. The presence of a 7-methylguanosine cap at mRNA 5’ ends enhances ribosome recruitment through interactions with specific translation initiation factors that also bind to the PIC. This process, referred to as “cap-dependent initiation” (CDI), is thought to account for most translation initiation in metazoan genes.

In contrast to CDI, some eukaryotic virus mRNAs have complex RNA structures that directly recruit ribosomes to internal start codons, termed Internal Ribosome Entry Sites (IRESes). First identified in Poliovirus and Encephalomyocarditis virus (EMCV)^3,4^, IRESes have been reported for hundreds of viruses, and can be classified into six major different types based on their RNA structural characteristics and their requirements for eukaryotic initiation factors (eIFs)^5,6^. IRES Types I, II, and III each fold into characteristic structures and require different sets of eIFs, including eIF4G. Type IV IRESes, exemplified by the human Hepatitis C Virus IRES, directly bind to the 40S ribosomal subunit and require only eIF2 and eIF3. Type V IRESes require both eIF4G and DHX29^7^. Unlike Types I-V, Type VI IRES elements directly bind to 80S ribosomes, and forgo the requirement for standard initiation factors. These elements fold into triple pseudoknot structures that mimic a hybrid elongation state^8^. However, only some viruses use IRES elements, as they are one of several strategies by which viruses express multiple protein coding sequences from single mRNA transcripts^9^.

Shortly after their discovery in viruses, researchers reported analogous IRES-like elements in mammalian genes, generally termed “cellular IRESes”^10^. Cellular IRES candidates were first proposed based on their apparent resistance to translation suppression during stress, due to their presence in dense polysome fractions on sucrose gradients^11,12^. To further study such elements, researchers developed assays intended to place cellular IRES candidates upstream of reporters expected to prevent CDI. Cellular IRES-like activities have been experimentally studied by testing their ability to drive translation in plasmid reporters that place the test sequence either downstream of another ORF (bicistronic) or in a context to make circular RNA. However, both plasmid reporter systems are known to produce false-positive IRES signals due to promoter activity and cryptic splicing in test elements that can create monocistronic transcripts translated by CDI^13–20,22^. As such, though hundreds of cellular IRES candidates have been nominated, it remains unclear which, if any, of these sequences are bonafide IRES elements.

The identification and study of viral and cellular IRES activities is highly important. In addition to understanding basic mechanisms of gene regulation, which could lead to antiviral drug targets, IRES elements are very useful for biotechnological applications, including RNA therapeutics^21^. Two previous studies screened thousands of human and viral gene fragments and reported IRES activities. The first study used a bicistronic reporter structure with candidate IRESes inserted between mRFP and eGFP in lentiviruses. Cells transduced with the lentivirus library were sorted by FACS to identify IRES candidates with relatively high eGFP expression, compared to mRFP^23^. A second study screened the same IRES candidates using a plasmid designed to make circRNA by backsplicing, in a library designed to transfect individual cells with single plasmids^24^. Each study reported thousands of seemingly functional IRES elements in human and viral genes, including important human pathogens like CMV, KSHV, HPV, Ebola virus, West Nile Virus, and HIV. Yet, there was no correlation between the measured strength of candidate IRES activities in bicistronic and circRNA contexts^24^. Although false-positive IRES calls are known to occur with both bicistronic and backsplicing reporter assays, none of the positive IRES candidates were validated using direct RNA transfection. Furthermore, these DNA-based MPRAs could not compare the relative magnitudes of CDI and IRES initiation from 5’ UTR sequences, and did not have the temporal resolution needed to assay potential IRES activity under stress. Thus, there is a strong need for a high-throughput IRES MPRA that does not rely on *in cellulo* transcription of stable fluorescent proteins.

Here, we present IRES-<u>Tr</u>anslating <u>A</u>ffinity <u>P</u>rotein <u>Pr</u>ofiling (IRES-TrAPPr), a novel RNA-based high-throughput platform for IRES discovery. IRES-TrAPPr uses affinity purification of nascent proteins to directly compare CDI and IRES activity from candidate IRES elements. Using stringent positive and negative controls, we show this method is highly specific and corresponds to orthogonal expression assays with high precision. IRES-TrAPPr quantitatively measures differences in IRES activities, as shown by structure function analysis of the IAPV and HCV IRESes. We used IRES-TrAPPr to compare the activities of hundreds of Type IV and Type VI viral IRESes in human cells, which identified highly active elements that drive similar rates of IRES and CDI translation. Analysis of these IRESes suggests that RNA structural stability is a key factor contributing to viral IRES tropism. Finally, we found negligible IRES activity from over a thousand cellular and viral IRES elements previously identified in bicistronic and circRNA plasmid MPRA screens. Our findings establish IRES-TrAPPr as a sensitive and accurate method for high-throughput IRES screening and structure / function analyses.

## Results

### An RNA-based MPRA to compare IRES activity and cap-dependent initiation

To avoid well-documented IRES false-positive artifacts common to plasmid-based IRES assays, we developed IRES-TrAPPr, a new MPRA inspired by NaP-TRAP^25^. In NaP-TRAP, a library of 5’ UTR variants are cloned upstream of a reporter protein containing an N-terminal affinity tag and a C-terminal degron, allowing for enrichment of nascently translating mRNAs.

IRES-TrAPPr places 5’ UTR variants upstream of two separate mRNA reporter libraries - a CDI reporter library containing a 5’ m7G cap (G-cap) and an IRES reporter library initiating with an adenosine followed by a highly stable stemloop structure (A-cap) to prevent CDI initiation (Figure 1A). A short barcode sequence is included in A-cap 5’ UTRs to identify A-cap and G-cap mRNAs by sequencing (see methods). A-cap and G-cap libraries are co-transfected into tissue culture cells, and nascently translating mRNAs are immunoprecipitated, washed at 37℃ to remove non-specific binding, and eluted by TEV protease cleavage. These translating mRNAs are then converted to cDNA and subjected to RNA-sequencing to quantitate individual 5’ UTR variants from A-cap and G-cap RNAs. The translation efficiency of each 5’ UTR construct is then estimated as the log ratio of the levels in the immunoprecipitated pool vs the input pool.

**Figure 1:**
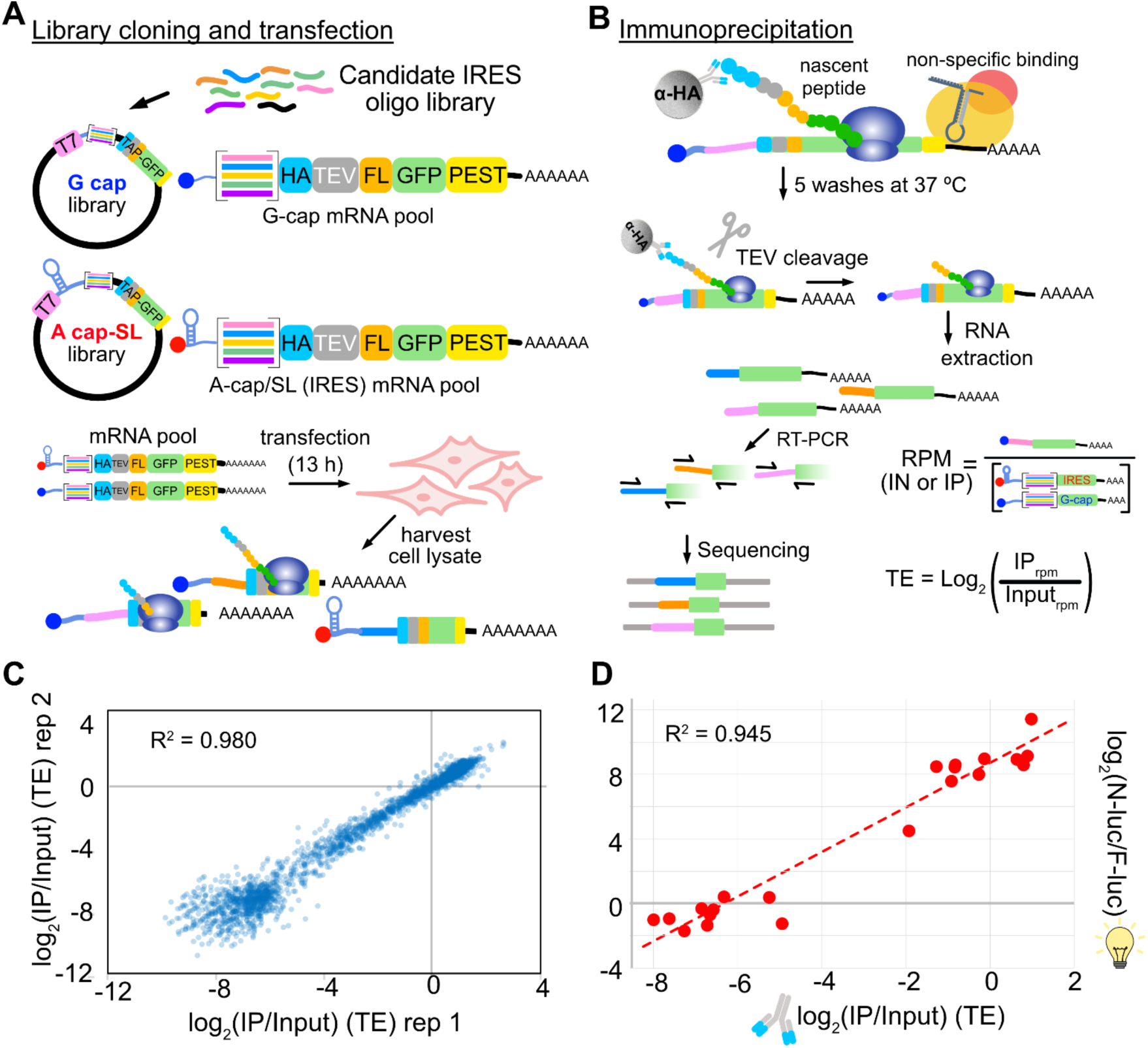
IRES-TrAPPr quantitatively compares IRES- and cap-dependent translation. **A)** Candidate IRES sequences are cloned into the 5’ UTRs of cap-dependent (G-cap) and IRES (A-cap/SL) reporter plasmids designed for immunoprecipitation of nascently translating mRNAs. Cap-dependent (G-cap) reporter RNAs are transcribed using a 5’ m?G cap, while IRES reporter RNAs incorporate a 5’ adenosine cap (A-cap) immediately followed by a stable stemloop (SL). IRES and G-cap mRNAs are cotransfected into HEK293T cells and cytosolic lysates are harvested. **B)** Nascently translating RNAs are immunoprecipitated using anti-HA beads, washed at 37°C to remove non-translating contaminant RNAs and eluted from the beads by TEV protease cleavage. Eluted RNAs are extracted and sequencing libraries are generated by RT-PCR using shared universal priming sites. The resulting libraries are sequenced to identify and quantify the Reads per million (RPM) for each candidate IRES and G-cap construct. The translation efficiency (TE) is calculated as the log_2_ ratio of RPM values for IP and Input **(IN)** samples. **C)** Example correlation of TE values for IRES and G-cap RNAs from two biological replicates. **D)** Validation of IRES-TrAPPr TE values. Twenty-two IRES and G-cap reporter constructs were cloned upstream of nanoluciferase reporters and individually tested by co-transfecting with a G-capped F-luciferase mRNA control reporter in HEK293T cells.

We designed an initial library of nearly two thousand candidate IRES sequences as IRES-TrAPPr reporters. These included negative control sequences that have negligible activity in both circRNA plasmid and RNA transfection assays^22^, and positive control IRESes from the HCV, IAPV, and CrPV viruses. For IAPV, the library contained 230 sequence and structure variants. Our library also included 93.5% of human and 89.7% of viral IRES candidates in IRESbase^26^, which covered all of the novel human and viral 5’ UTR IRES elements reported in the DNA-based bicistronic MPRA study^23^ and many additional previously reported IRESes. We also included 401 reportedly highly active IRESes from full length human genes, including genes whose mRNAs remain polysomal in response to stress^23^. In addition, we included the 5’ UTRs of Zika and Dengue virus, which have had conflicting reports of IRES activity^27–30^. Using this library, we performed IRES-TrAPPr in HEK293T cells with three biological replicates, each of which had two technical replicates for sequencing library preparation. IRES-TrAPPr translation efficiency estimates spanned a 1,000-fold range, were highly reproducible (Figure 1B; Figure S1) and were validated by independent transfection of twenty-one IRES-candidate mRNAs cloned upstream of nanoluciferase (Figure 1C). These results show that IRES-TrAPPr provides accurate reproducible estimates of IRES and CDI translation activity.

### Dissecting IRES structure / function relationships

Viral IRES elements recruit ribosomes by folding into specific, sometimes elaborate structures. To evaluate the utility of IRES-TrAPPr for studying viral IRES structure / function relationships, we tested the impact of mutations in two well-characterized IRES elements on their ability to recruit ribosomes for translation. The IAPV IRES secondary structure includes multiple paired helices and pseudoknots (Figure 2A, Table S1)^31^. We used scanning mutagenesis to quantify the impacts of trinucleotide mutations across the IAPV IRES sequence. Mutations in known functional regions decreased IRES activity up to 80-fold. Mutations that disrupted known secondary structural interactions and pseudoknots were most impactful. Like many Type VI IRESes from dicistroviridae, IAPV is known to drive translation in two reading frames - the first of which initiates at a GGC “Gly” codon and the second which initiates at a GCG “Ala” codon. Notably, our scanning mutagenesis identified mutations in IAPV that increased IRES translation in the first reading frame 2-fold, potentially by reducing translation initiation in the +1 frame. To investigate this, we compared +1 frame translation from the IAPV IRES in luciferase assays and found the mutations that increased frame 0 TE in IRES-TrAPPr (m203 and m146) decreased +1 frame translation (Figure S2). In summary, IRES-TrAPPr can be used to quantify a wide range of IRES variants, providing structural and functional dissection of reading-frame specific IRES activities.

**Figure 2.**
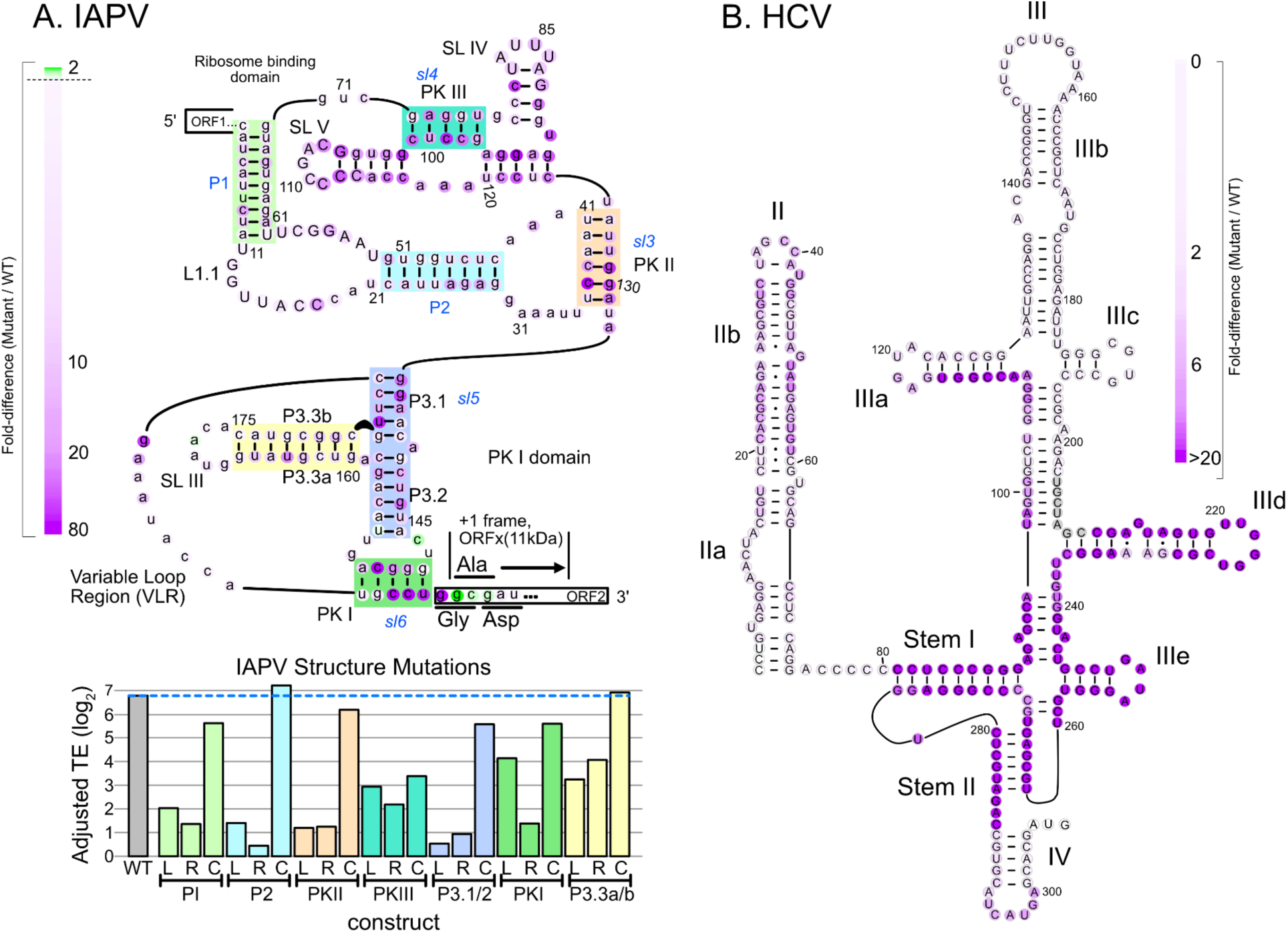
Dissection of IRES structure *I* function relationships using IRES-TrAPPr. **A)** Structure/ function analysis of the IAPV IRES. (Top) The translation efficiencies of IRES constructs with three nucleotides adenosine mutations (AAA) in one nucleotide steps (1-4, 2-5, 3-6, etc) were compared to wildtype IAPV. The center nucleotide (nt 2 for 1-3, AAA) was colored to reflect the fold decrease for each mutation across an 80-fold range of activity. Mutations to the +1 frame Ala codon increased +0 frame translation two fold (green nucleotides). (Bottom) The bar graph indicates the relative translation efficiencies of IAPV IRES variants harboring mutations in known basepairing helices, shown as colored boxes in the secondary structure diagram. For each region, all of the nucleotides on one side of the helix were mutated to ablate base-pairing potential on the left (L), or right (R) side of the helix. Compensatory (C) mutations restoring basepairing potential for each helix were created by combining Land R mutations. Except for Pseudoknot 111, all compensatory mutations increased translation efficiency as compared to the Land R mutations, validating the known helices in the IAPV IRES structure. **B)** Scanning mutagenesis across the HGV IRES identifies sequences critical for IRES function. IRES constructs were cloned in which four-nucleotide segments were mutated to remove basepairing potential (A<-= C; G <-= **T)** in three nucleotide steps (1-4, 4-7, 7-10, etc.). The mutant IRES mRNA reporter pool was transformed into HEK293T cells and assayed by IRES-TrAPPr. The IRES translation efficiency for each mutant was compared to wildtype to calculate the fold-decrease caused by each mutation across a 50-fold range. Nucleotides were oolored by the fold-change effect.

We further reasoned that IRES-TrAPPr could be used to test the importance of specific IRES structural elements. We evaluated this by testing a series of “left” (L), “right” (R), and “compensatory” (C) mutations to each base-paired region and pseudoknot in the IAPV IRES by converting all nucleotides on one (L and R) or both (C) sides to their complementary base. The L and R mutations disrupted IRES activity from eight to eighty-fold and, with the exception of pseudoknot III, compensatory mutations largely restored IRES activity. Finally, we constructed a similar scanning mutagenesis library of mutations in the Type IV HCV IRES. Similar to our results with IAPV, this verified the known structure^33,34^ of the HCV IRES and identified the core functional regions of the HCV IRES involving stems I, II, IIIa, IIId, and IIIe (Figure 2B; Table S2), including the critical “GGG” loop in stem IIId that directly contacts 18S ribosomal RNA ES7^32^. Together these results demonstrate that IRES-TrAPPr identifies sequences and structures that are critical for IRES activity.

### Determining the relative activities of hundreds of viral IRESes in human tissue culture cells

Phylogenetic studies have identified hundreds of viral IRES candidates with sequence and structural similarity to the IAPV and HCV IRESes^5,6,35,36^. These IRESes were found in virus sequences isolated from a wide array of vertebrate *picornaviridae* (Type IV) and invertebrate *dicistroviridae* (Type VI) hosts by searching for both sequence and RNA structure homology, and some have been shown to be active in human cells and *in vitro* translation systems. Having established the sensitivity and specificity of IRES-TrAPPr, we next used the assay to evaluate the relative strengths of such elements in driving translation in human cells. We tested a library containing 37 Type IV and 182 Type VI candidate IRESes. Using the TE of the CrPV IRES as a lower boundary for “active” IRES classification, we identified 43 active Type VI and 24 active Type IV IRESes (Figure 3A; Table S3). Thus, most Type IV IRESes drove cap-independent translation at a detectable level in HEK293T cells, while most Type VI IRESes did not. However, several Type VI IRESes endemic to insects had activities rivaling that of the human HCV IRES, three of which were validated in separate luciferase experiments (Figure 3D). We also compared IRES activity to CDI translation from these elements. This is important, as candidate IRESes that exhibit low rates of IRES activity, as compared to CDI, are unlikely to be biologically relevant IRESes^37^. In addition, mis-mapping of even a few sequence reads can create the appearance of weak IRES activity. Using CrPV as a lower bound for activity, we found active IRESes drove 46% as much IRES translation as CDI on average, while inactive IRESes drove only 4% as much translation (Figure 3B). These results establish IRES-TrAPPr as a powerful method for identifying bonafide IRES activities in human cells.

**Figure 3.**
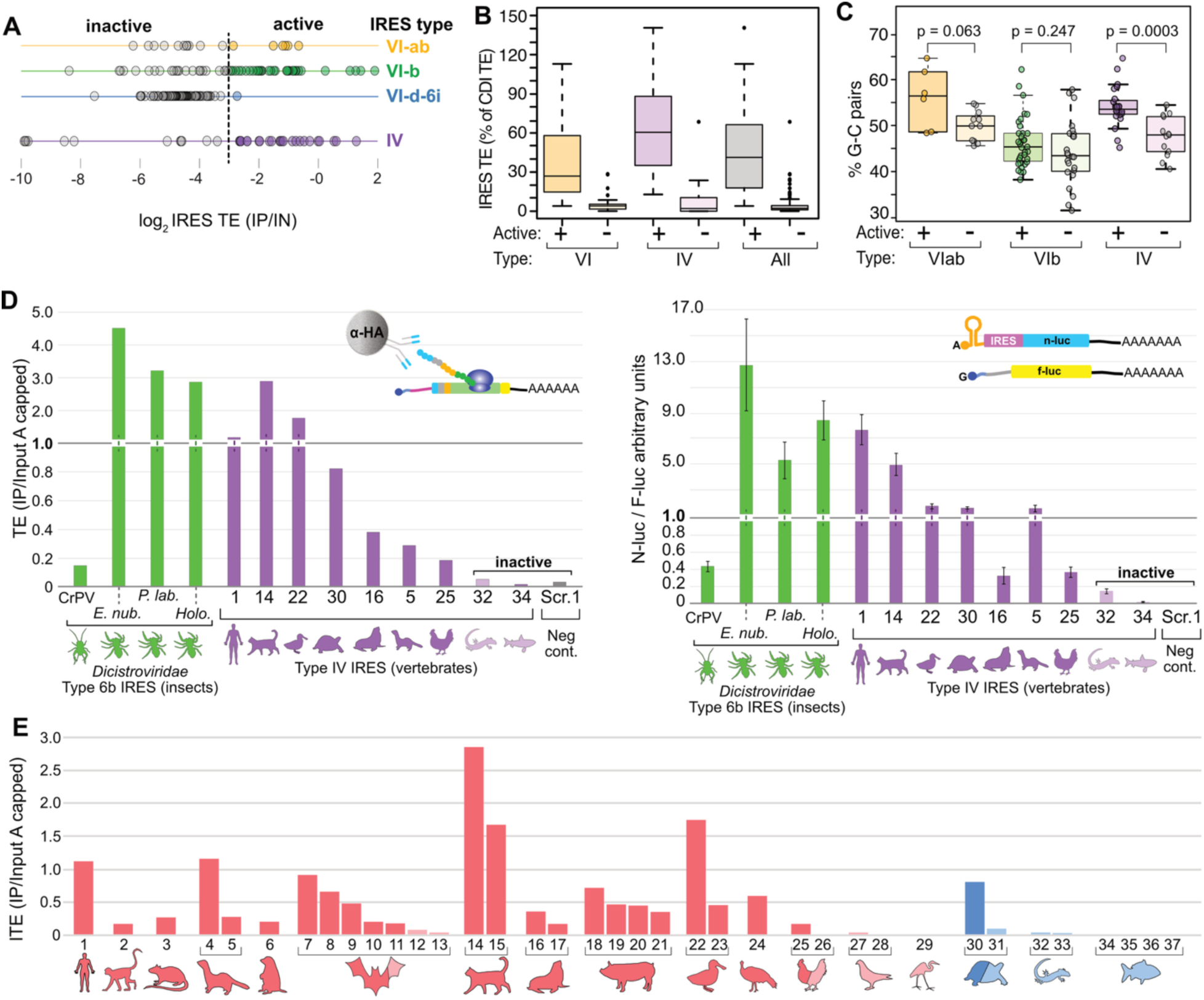
Characteristics of Type IV and VI IRESes with high activity in human cells. **A)** Line graphs show the relative translation efficiencies (TEs) of Type VI (yellow, green, blue) and Type IV IRESes in HEK293T cells. The dashed line indicates the activity level of the CrPV IRES. **B)** Boxplots of IRES TEs as a percentage of CDI TEs from the same candidate IRESes. **C)** Boxplots comparing the percentage of all basepairs that are strong G-C pairs for “active” and “inactive” Type VI and Type IV IRESes. IRESes that are active in human cells have higher frequencies of G-C pairs, compared to IRESes with no activity in human tissue culture. **D)** Barplot showing TE values (left) and luciferase validation assays (right) for highly active Type VI and Type IV IRESes. Three Type Vl-b IRESes isolated from arachnid hosts drive translation comparable to the HCV IRES in HEK293T cells. Type IV IRESes are numbered as described below. **E)** Barplot shows IRES-TrAPPr TE values for Type IV IRESes isolated from endotherm (left, red) and ectotherm (right, blue) host species. IRESes isolated from ectotherm hosts are generally inactive in human cells. 1-HCV, 2-Simian sapelovirus, 3-Manhattan virus, 4-Ferret parechovirus, 5-Ferret kobuvirus, 6-Marmot mosavirus, 7-Bat picornavirus 1, 8-Bat crohivirus, 9-Bat picornavirus 2,10-Bat hepatovirus, 11-Bat kunsagivirus, 12-*la io* bat picornavirus, 13-Diresapivirus B1, 14-Feline picornavirus, 15-Feline sakobuvirus A, 16-California sea lion sapelovirus 1, 17-Seal picornavirus 1, 18-Porcine sapelovirus 1, 19-Porcine teschovirus, 20-Senecavirus A, 21-Swine pasivirus 1, 22-Duck hepatitis virus **1,** 23-Pink-eared duck picornavirus, 24-Turkey hepatovirus 1, 25-Avian encephalomyelitis virus, 26-Chicken megrivirus, 27-Pigeon picornavirus B, 28-Pigeon mesivirus, 29-Grusopivirus A1, 30-Tortoise rafivirus A1, 31-Pemapivirus A1, 32-Tropivirus A1, 33-Guangxi Chinese lizard picornavirus 2, 34-Fathead minnow picornavirus, 35-Bluegill picornavirus, 36-Symapivirus A1, 37-Carp picornavirus

Type VI IRESes have been classified into a number of subtypes based on their length and predicted structural features^6^. Notably, the most active Type VI IRESes come from the VIab and VIb subtypes. However, many of these IRES candidates were inactive in our system. As these elements originated in viruses that infect invertebrates, it is possible that some may be poorly suited to fold into active IRES structures at the relatively high temperature in which human cells grow. To investigate this, we compared the percentage of strong GC base pairs in the predicted structures of active and inactive Type VIab and VIb IRESes. Active IRES elements had more GC pairs than inactive ones, though this only approached significance for Type VIab IRES elements and was not statistically significant for Type VIb (Figure 3C). Strikingly, we found a stronger trend among active and inactive Type IV IRES elements from vertebrate hosts, suggesting that structural stability contributes to viral IRES adaptation to host cells.

Furthermore, among Type IV IRESes, we found that IRESes isolated from endothermic hosts were generally more active in human cells than those isolated from ectothermic hosts (Figure 3E). Indeed, the only IRES element isolated from an ectothermic host that had activity in human cells came from a tropical tortoise, while all IRESes isolated from fish, turtles, and lizards were inactive. Together, these results suggest that Type IV IRESes from viruses infecting warm blooded animals may have evolved to be more structurally stable at higher temperatures.

### Evaluating viral and cellular IRES candidates nominated by DNA-based assays

We next used IRES-TrAPPr to evaluate the activity of 499 viral and 646 cellular IRES candidates previously nominated using DNA-based assays and collected in a public database^26^. This included all of the human and viral 5’ UTR IRESes reported to function in human cells in the original IRES MPRA study^23^ and IRESes reported from the 5’ UTRs of Dengue and Zika viruses. Surprisingly, only two viral IRESes from IRESbase were categorized as “active” by IRES-TrAPPr (Figure 4A, Table S4), both of which are Type VI IRESes. Overall, the IRES TE of nearly all of the IRESbase viral IRES candidates we tested was very low compared to CDI translation (Figure 4B). In addition, we found no evidence of IRES activity from the Dengue and Zika virus 5’ UTRs. Instead, the Zika and Dengue 5’ UTRs promoted extremely efficient translation by CDI. Independent validation by transfection of mRNA luciferase reporters verified these results (Figure S3). These findings suggest that the vast majority of short viral IRES elements proposed using DNA-based assays, including MPRAs, are false positives, potentially resulting from artifacts that occur in plasmid reporter systems. This underscores the value of using RNA-based reporter systems to screen for IRES activity.

**Figure 4.**
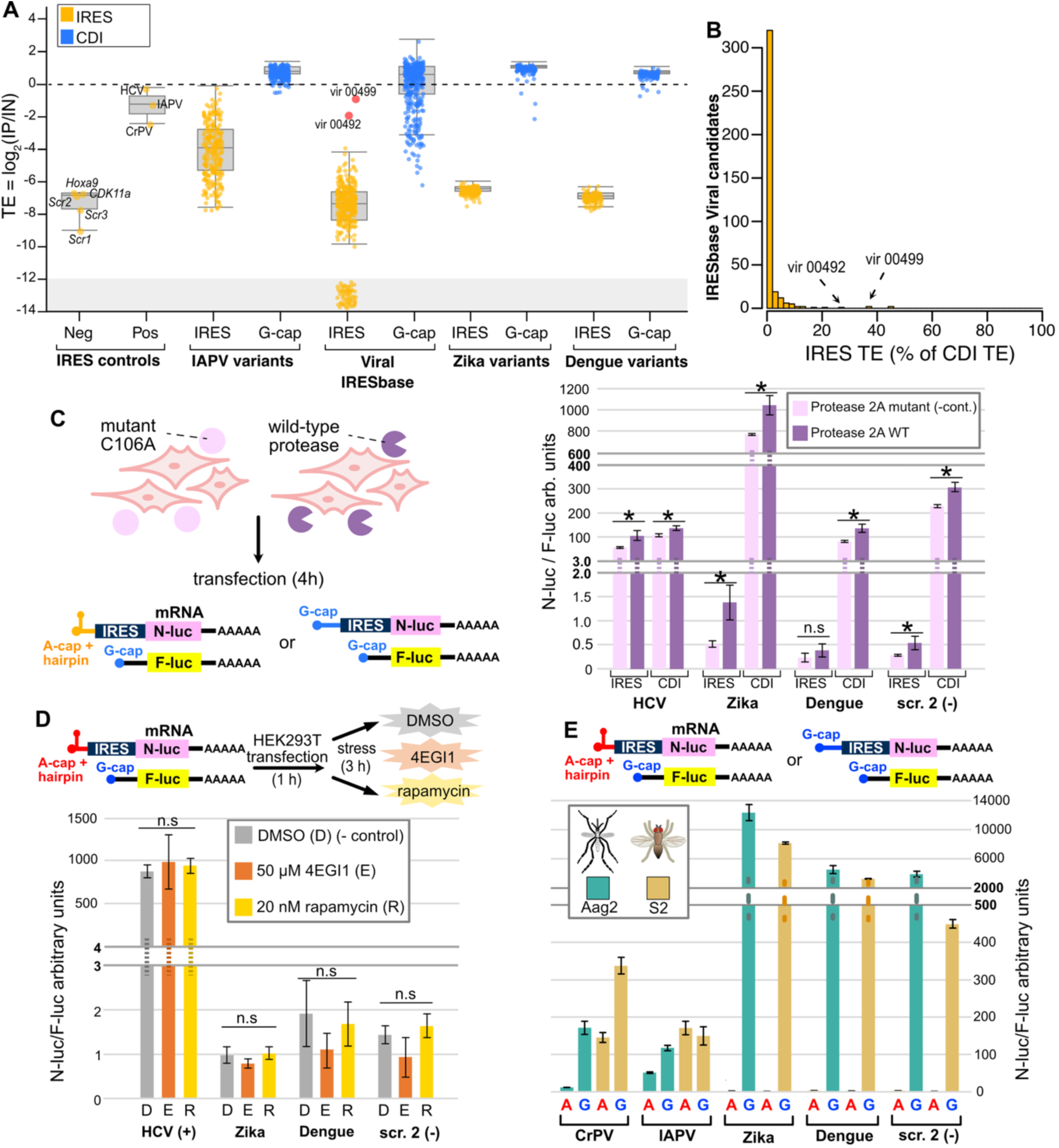
IRES-TrAPPr finds previously reported short viral IRESes have negligible IRES activity. **A)** Boxplots show the TEs for IRES and G-cap TRAP reporters. Positive control IRES elements have much higher IRES activity than negative controls. The IAPV variants from Figure 2 depict the range of IRES activities resulting from mutations to the IRES structure. Short viral IRESes from IRESbase, primarily from Weingarten-Gabbay et al., were almost universally negative in this assay, as were candidate IRES elements from the Zika and Dengue virus 5’ UTRs. Log_2_ TE values less than -12 represent samples that had zero reads in the IP and were assigned pseudocounts. **B)** Histogram depicts the IRES TE for IRESbase viral IRESes as a percentage of their CDI translation **C)** Diagram shows expression of wildtype and inactive C106A mutant Rhinovirus 2A protease, which cleaves elF2A and other targets. Contrary to previous studies, HRV 2A protease did not induce appreciable IRES activity from Zika and Dengue virus 5’ UTRs. Cap-dependent translation from these 5’ UTRs was much more efficient in the presence of both wildtype and inactive C106A mutant 2A protease. **D)** Translational stress with elF4EGl1 inhibitor and rapamycin failed to induce IRES activity from Zika and Dengue 5’ UTRs. **E)** The Zika and Dengue 5’ UTRs also had negligible IRES activity when transfected into the mosquito *Aedes aegypti* (Aag2) and *Drosophila* (S2) tissue culture cell lines.

Previous work reported that Dengue and Zika virus 5’ UTRs drove cap-independent translation only under conditions in which CDI was suppressed by stress or viral infection^28–30^. For example, it was reported that Human Rhinovirus (HRV) 2A protease, which cleaves eIF4G, induces IRES-like translation from the Dengue virus 5’ UTR^30^. To further investigate, we transfected cells with wild type and mutant HRV 2A protease and compared IRES and CDI translation using luciferase reporter constructs. Contrary to a previous report, we found negligible IRES activity from Zika and Dengue 5’ UTRs after stress. Indeed, both CDI and IRES-mediated translation appear to be upregulated to similar extents after HRV 2A expression (Figure 4C), which largely reflects reduced translation of both flavivirus Nluc and control Fluc luciferase. Similarly, we found chemical inhibition of CDI did not induce IRES activity from the Zika and Dengue Virus 5’ UTRs (Figure 4D and S3). Finally, because these viruses are carried by insect vectors, we tested their 5’ UTRs in mosquito (Aag2) and fruit fly (S2) cell lines in the event that IRES translation is used primarily in insect hosts. While IAPV and CrPV IRES elements were active in these cell lines, no IRES activity was found for Zika or Dengue 5’ UTRs (Figure 4E). These results show that the Zika and Dengue 5’ UTRs have negligible IRES activity.

We next evaluated the activity of potential cellular IRESes previously nominated using DNA-based reporter assays. Similar to our findings with candidate viral IRESes, we found negligible activity from cellular IRES candidates, including all of the candidate 5’ UTR IRESes reported in the original bicistronic MPRA screen and candidates from genes reported to be insensitive to CDI inhibition during stress (Figure 5A, Table S5). In concordance with this interpretation, we further tested four of the most active candidates using luciferase mRNA transfection and found IRES activity similar to that of the random sequence negative control (Figure 5B). We also found nearly all cellular IRES candidates drove very little translation by IRES activity, relative to CDI (Figure 5C). These results show candidate cellular IRESes nominated by DNA-based bicistronic reporter systems have negligible IRES activity in our RNA-based IRES-TrAPPr MPRA.

**Figure 5.**
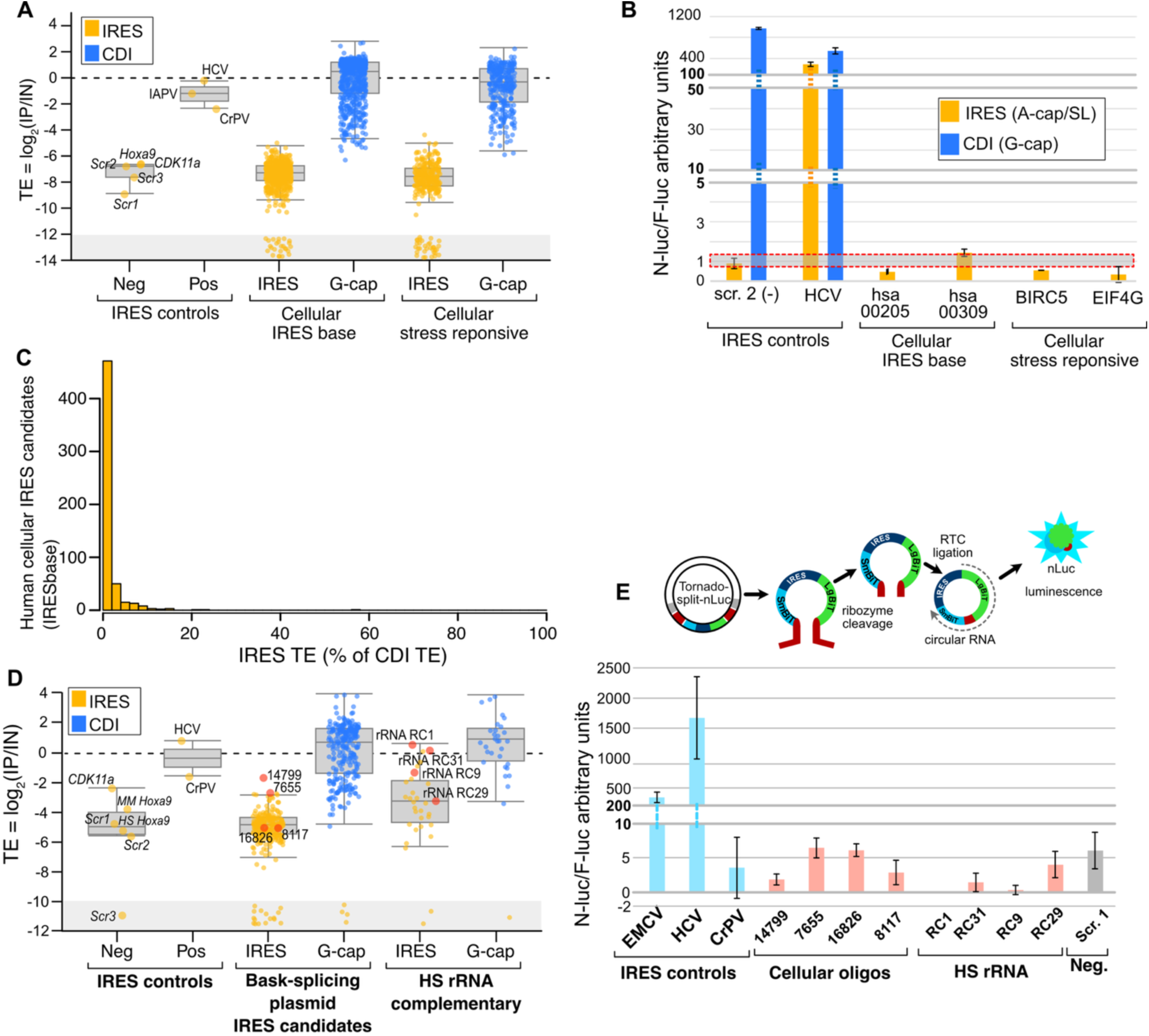
Previously reported cellular IRES candidates have negligible IRES activity. **A)** Boxplot shows the TEs for IRES and G-cap RNA reporters measured by IRES-TrAPPr. Candidate cellular IRESes (IRESbase) have negligible IRES activity, similar to the negative controls, regardless of their origin in stress response genes (right) and much higher translation as G-capped RNAs (blue). **B)** RNA transfection of nanoluciferase reporters validates IRES-TrAPPr results for cellular IRES candidates. G-capped RNAs were not tested for the cellular IRES candidates in this experiment **C)** Histogram depicts the distribution of IRES **TE** as a percentage of CDI **TE** from the same sequences from cellular IRES candidates hosted on IRESbase. Nearly all candidate cellular IRES elements drove CDI at least 20-fold more than IRES initiation (5% IRES or less). **D)** Boxplots depict the TEs of candidate cellular IRESes that had the highest activities in backsplicing circRNA reporter. **E)** Candidate IRESes previously reported from a back-splicing plasmid screen (red dots from **D)** were tested using the Tornado plasmid in HEK293T cells to assay potential IRES activity in a circRNA context. None of the candidates had IRES activity in excess of the negative control. Note that CrPV IRES had poor activity in this plasmid.

We separately tested 265 cellular IRES candidates reported to have the highest IRES activity in an MPRA screen that used back-splicing circRNA plasmids^24^. While such plasmids have been shown to create false-positive IRES signals due to *trans*-splicing^15–17,22^, we reasoned the most highly active candidates may be more likely to include bonafide functional IRESes.

Nevertheless, these circRNA IRES candidates also had negligible activity in our assay (Figure 5D, Table S6). A few candidates appeared to have IRES activity, but these could not be validated using the Tornado circRNA plasmid system, which minimizes false-positive artifacts^19,38^ (Figure 5E). Finally, we tested a tiled array of 31 sequences complementary to the 18S rRNA as such elements were also reported to have IRES activity in two previous DNA-based IRES MPRA studies^39,40^. While several of these sequences were enriched by immunoprecipitation (Figure 5D), we were unable to validate their activity using the Tornado plasmid circRNA system (Figure 5E). Together these results indicate that putative cellular IRES elements nominated using DNA-based MPRAs have negligible activity in the RNA-based IRES-TrAPPr system. As DNA-based IRES assays are known to create artifactual IRES signals, it is likely that these putative IRESes are false-positives.

### Comparing the effects of cellular stress on IRES activity and cap-dependent initiation with IRES-TrAPPr

Protein synthesis by CDI is suppressed in response to acute cellular stress^41^. Because of this, mRNAs that are polysome associated during stress have been considered candidates for IRES activity^10,37^. Indeed, many genes reported to harbor candidate cellular IRESes were initially studied due to their apparent resistance to translation inhibition under stress^42^.

Furthermore, a major function of viral IRESes is to direct the synthesis of viral proteins during infection, when cellular translation is suppressed. Because it measures nascent translation, IRES-TrAPPr has high temporal resolution, creating an ideal platform to examine the effects of cellular stress on IRES activity. To investigate this, we transfected cells with our IRES-TrAPPr library carrying candidate viral and cellular IRESes, treated cells with thapsigargin or DMSO (negative control) for three hours, and prepared cellular extracts for IRES-TrAPPr. To compare translation between treated and control cells, we created a reference extract from unstressed cells transfected with three serially diluted spike-in control reporters. The spike-in control lysate was added to a final of 1% of each of the treated and control extracts before preparing and sequencing IRES-TrAPPr libraries (Figure 6A). Polysome analysis showed that thapsigargin treatment results in a substantial increase in monosome signal, relative to polysomes, consistent with reduced translation initiation (Figure 6B, Table S7). Our TE estimates from IRES-TrAPPr revealed that CDI reporters had an ∼40% decrease in translation efficiency in response to thapsigargin treatment (G-cap Figure 6C), consistent with inhibition of cap-dependent translation. At the same time, our positive control IRES sequences showed increased translation efficiency in response to stress, as did our IAPV variant library and the two active Type VI viral IRESes from IRESbase. Consequently, the IRES TE as a percentage of CDI TE increases strongly for bonafide viral IRESes in response to stress. In contrast, IRES activity from candidate viral and cellular IRES elements nominated by DNA-based assays was unchanged in response to stress. These results demonstrate that IRES-TrAPPr can detect stress-dependent changes in CDI and IRES activity, and that the large number of putative short cellular and viral IRESes nominated using previous methods are not induced by cellular translational stress.

**Figure 6.**
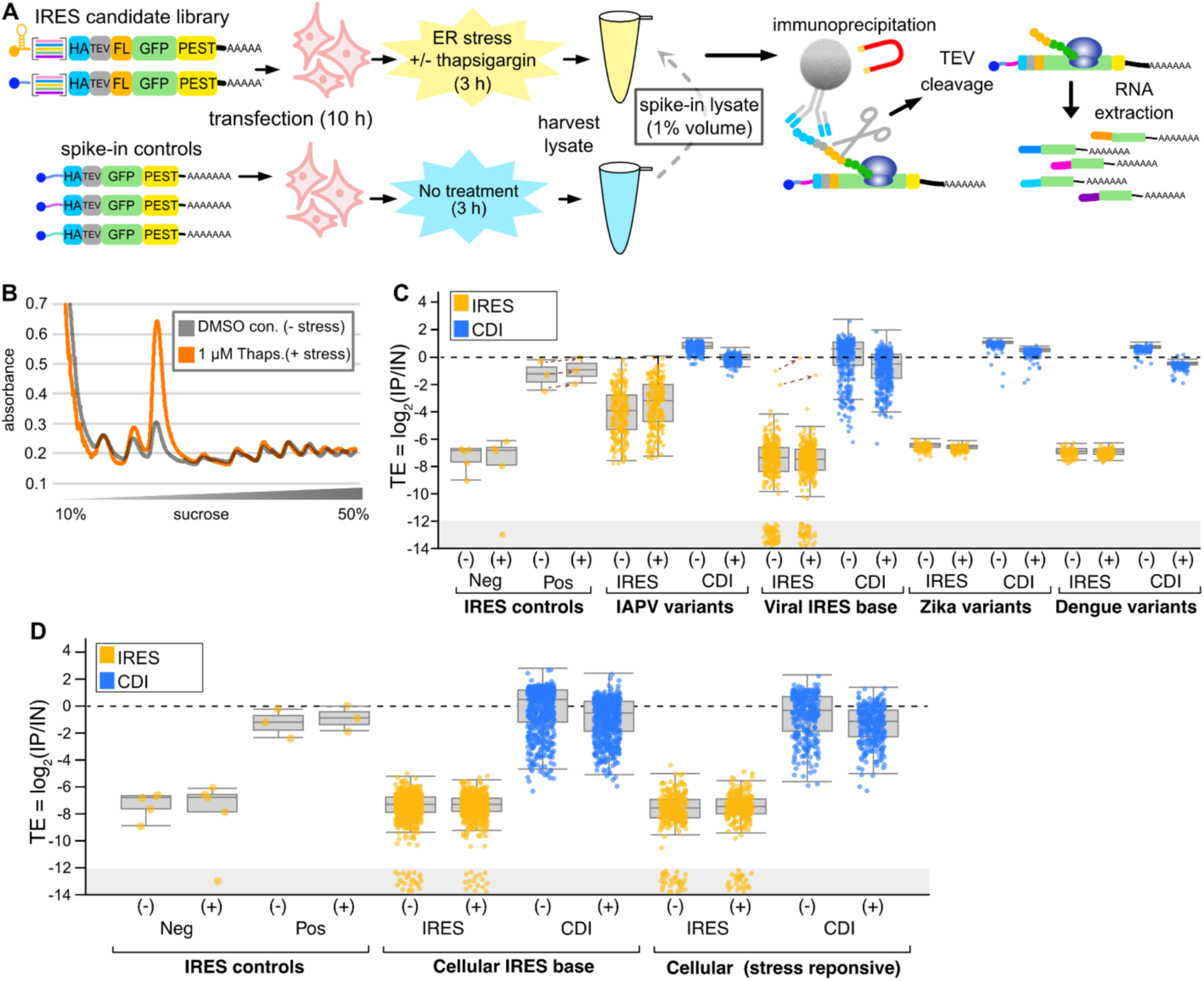
ER translational stress enhances IRES activity for bonafide viral IRESes, but has no effect on candidate cellular IRESes. **A)** Diagram depicting IRES-TrAPPr stress-response assay. The candidate IRES library is transfected into tissue culture cells for 10 hours, with and without a 3 hour exposure to 1 uM thapsigargin, and polysome lysates are harvested. Separately, three G-cap spike-in control RNAs, serially diluted 10-fold, are transfected into tissue culture cells and polysome lysates are prepared. The Spike-in control lysate is added to each experimental lysate to 1% final volume and mixed before subjection to the IRES-TrAPPr protocol. **B)** UV absorbance trace showing the effects of thapsigargin treatment on polysomes and monosomes. **C)** Boxplot depicts the spike-in adjusted translation efficiencies before(-) and after(+) ER stress. Viral IRES elements, including positive controls, IAPV variants, and the two functional viral IRESbase IRESes, had increased TEs after stress (dashed arrows), while cap-dependent translation was reduced. **D)** Candidate cellular IRESes showed no apparent increase in translation under stress, regardless of whether or not the IRES candidates originated from stress-response genes.

## Discussion

Here, we present IRES-TrAPPr, a novel RNA-based massively parallel reporter assay that quantitatively compares IRES activity to translation by cap-dependent initiation. IRES-TrAPPr provides many advantages for studying IRES activity compared to current state-of-the-art methods. Using a range of well-characterized viral IRESes as positive controls, and non-IRES sequences as negative controls, we demonstrate that IRES-TrAPPr is both highly specific and highly precise. Our scanning mutagenesis experiments validate IRES-TrAPPr, and illustrate its sensitivity and utility for RNA structure / function studies. We expect this will be very useful for studies of new IRESes and their functional structures, given the explosive increase in viruses that have been identified through the Global Virome Project^43^. Pinpointing nucleotides important for viral IRES function may also aid in the development of antiviral compounds targeting IRESes^44,45^. In addition, our scanning mutagenesis resulted in IAPV and HCV IRES variants that induce a 1,000-fold range of translation. These variants could be particularly beneficial for biotechnology applications as they provide specific tunable levels of expression from multicistronic synthetic mRNAs.

Our method is also ideal for identifying IRES elements that function in human cells. We quantified translation from over two hundred Type IV and Type VI IRESes in human cells and identified sixty-seven that drove at least as much translation as the CrPV IRES. The predicted structures of active IRES elements from both classes have a higher percentage of strong (GC) base-pairs than inactive IRESes, a trend that was strongly significant for Type IV IRESes. This suggests that some of these IRESes may have evolved to be structurally stable at higher host body temperatures. IRES elements isolated from ectothermic hosts have not evolved under selection for strong structural stability. Notably, all the Type IV IRESes from fish, lizard, and turtle viruses were inactive in mammalian cells, presumably due to misfolding of the IRES structures at higher temperatures. Indeed, the only functional Type IV IRES from an ectothermic host was isolated from a tortoise endemic to the tropical island of Sulawesi. We also found that two of the seven Type IV IRESes isolated from bats had very low activity in HEK293T cells. It is possible that these viruses replicate during hibernation, when bat body temperatures drop dramatically. Our screen also identified Type VI IRESes from arachnid hosts with activities in human cells rivaling or exceeding that of the HCV IRES. These high-activity IRESes may be particularly useful for synthetic biology, as Type VI IRESes have the unique ability to initiate translation in two reading frames. Notably, many Type IV and Type VI IRES candidates were not functional in HEK293T cells. These may require different cellular environments or cofactors to drive robust translation. Future work using IRES-TrAPPr in a variety of cell lines and species may identify unique requirements for the activity of specific viral IRESes.

In contrast to these positive IRES examples, IRES-TrAPPr also identified hundreds of inactive viral and cellular IRES candidates which were previously nominated in DNA-based high-throughput screens. Our results cast doubt on the often cited estimate that 10% of human genes have active IRES elements from the first high-throughput DNA-based IRES screen^23^. However, in hindsight, the design of the original screen may have precluded detection of bonafide IRESes. First, the technology of the time limited candidates to only 174 nucleotide fragments, which is much smaller than most IRESes from mammalian viruses. Indeed, none of the reported short candidate IRESes from human pathogens (HIV, CMV, KSHV, HPV, etc) were active in IRES-TrAPPr. Furthermore, the fragments screened in previous studies were designed by sliding window scanning with a 50-nucleotide step size. Consequently, most functional elements would have been cloned out-of-frame with the reporter protein. While these design constraints may have prevented detection of bonafide cellular IRESes, it is also possible that such elements are exceedingly rare in human genes as candidates routinely fail validation attempts using methods that resist false positives^13,22^. Candidate viral IRESes nominated by older studies were also entirely inactive in our assay, including reported IRESes from Zika and Dengue virus^27^ and sequences complementary to 18S rRNA^23,24,39,40^. However, it is still possible that new viral and cellular IRES elements remain undiscovered, including IRESes whose length exceeds the limitations of earlier studies and IRESes that require host-specific co-factors not present in our screen. IRES-TrAPPr provides a powerful platform to search for such activities.

Our results also raise important questions about public databases of IRES elements and computational modeling to predict IRES activity. Several databases are available and the candidate IRES elements listed therein have been used by multiple studies to develop machine learning models for IRES prediction, including IRESpy^46^, IRESpredictor^47^, DeepCIP^48^, DeepIRES^49^, IRESeek^50^, LAMAR^51^, and UTR-LM^52^. It is possible that the features used by these models to classify “IRES” vs. “non-IRES” reflect the mechanisms that drive false positives (e.g. promoters and splice sites), as ML algorithms can find features that discriminate between two groups even when the groups are mislabelled^53^. Current IRES databases are largely populated with candidate elements nominated by DNA-based screens, nearly all of which are inactive in IRES-TrAPPr. To our knowledge, none of the ML models trained on DNA-based IRES candidates have identified novel IRESes that function in RNA-based reporters. In contrast, several studies have found active IRESes by searching for sequence and structural homology to Type IV and Type VI IRESes^6,35,36^. Removing false-positive candidates from IRES databases may facilitate the development of better ML models that learn features of bonafide IRESes.

Defining a minimum “active” IRES threshold is an important consideration when screening candidate IRESes. 5’ UTRs that lack IRES elements may produce background levels of IRES reporter translation, but this signal is expected to be much lower than CDI translation from the same sequence^37,54^. By directly comparing IRES and CDI translation from the same sequences, IRES-TrAPPr allows researchers to evaluate the relative biological importance of apparent IRES activity. In doing so, we found that CDI was much more efficient than IRES initiation for viral and cellular IRES candidates nominated by DNA-based MPRAs (Figure 3B, 4B, and 5C). These inactive IRES candidates typically had very low reads in IP samples, potentially reflecting background noise in the assay. Indeed, we recovered rare A-cap/G-cap mismapped reads from controls that were added entirely as A-cap (scramble negatives) or G-cap (spike-in) transcripts. To account for such noise, we used the TE of the CrPV IRES as a minimum threshold for IRES activity. This element, originating in a cricket virus, has relatively weak IRES activity in human cells. We feel this is an appropriate threshold for identifying significant IRES activity, especially since the resulting “inactive” candidates have very low IRES TE as a percentage of CDI TE (Figure 3B). Furthermore, we were unable to validate multiple cellular IRES candidates whose apparent activity fell between the negative controls and CrPV.

Many viruses use IRES elements to direct translation during infection, when CDI translation is suppressed. It has long been suspected that human genes whose translation is resistant to stress may similarly use IRESes to drive translation during acute stress^37^. Because DNA-based IRES screens have low temporal resolution, their usefulness in studying IRES activity under stress is limited. By measuring nascent translation with spike-in controls, IRES-TrAPPr is perfectly suited to assay both CDI and IRES activity in response to cellular stress. We show that thapsigargin treatment both inhibits CDI and enhances IRES activity in HEK293T cells. Importantly, previously nominated candidate IRESes from genes whose translation appears to be resistant to stress also had no apparent increase in IRES activity after stress. This suggests other mechanisms may be responsible for translation of stress-resistant genes. For example, upstream Open Reading Frames (uORFs) are known to provide stress-resistant translation initiation in a number of contexts^41,55^. IRES-TrAPPr should facilitate future studies into IRES activities in response to a variety of stresses, including viral infection.

Although IRES-TrAPPr provides a highly accurate platform for quantifying and comparing IRES and CDI translation, there are several limitations to this method. First, as with all MPRA technology, IRES-TrAPPr is restricted by the technological limits of DNA oligo synthesis and high-throughput sequencing. Currently this allows inexpensive pools of IRES candidates up to 310 nt long. Many mammalian IRESes are longer (e.g. Poliovirus and EMCV) and require costly individual synthetic constructs. Illumina sequencing is also limited to 2 × 300 cycles at present, meaning full-length 5’ UTR sequencing has a practical limit of ∼ 500 nt. It is possible this limitation could be circumvented by instead sequencing barcodes that represent each IRES candidate. Finally, false-positive IRES activity, though rare, can still occur with sequences that are highly complementary to ribosomal RNA or other translating RNAs (Figure 5D). This is most likely due to Co-IP of non-translating reporter RNAs that base-pair to actively translated sequences^22^. Further optimization of wash times and buffers could potentially alleviate this issue, however validation of individual positive candidates by direct RNA transfection is still advisable.

Since the discovery of viral IRES elements nearly four decades ago, researchers have uncovered intricate RNA structures and molecular mechanisms viruses use to recruit ribosomes, while also searching for similar elements in other viruses and metazoan genes.

Indeed, recent studies using DNA-based MPRA systems have nominated thousands of candidate viral and cellular IRESes. However, DNA-based IRES assays are susceptible to false-positive artifacts caused by the presence of transcriptional promoters and cryptic splice sites in candidate IRES sequences^13,14,56–58^. RNA-based reporters are not subject to such issues and are thus considered a gold-standard for testing IRES activities^13,58^ but, thus far, have been limited to studying candidate elements on an individual basis. With the development of IRES-TrAPPr, we present a new RNA-based high-throughput platform for identifying functional IRESes and studying their structures. The many advantages of this system in accuracy, sensitivity, specificity, and temporal resolution will allow, for the first time, high confidence high-throughput research on cap-independent translation mechanisms.

## Methods

### Construction of GFP-Trap vectors

Vectors T7-GFP and T7HP-GFP(^22^) were used as backbones for library construction. The native GFP coding sequence was replaced with a modified GFP sequence containing a codon-optimized 3×HA–TEV–3×FLAG tag inserted 7 amino acids downstream of the start codon (referred to as GFP-Trap). To reduce GFP accumulation and increase the likelihood of capturing nascently translated proteins, a PEST degron was added downstream of GFP. The GFP-Trap sequence was synthesized as a gene fragment (Twist Bioscience, Table S8), digested with BglII and XbaI, and cloned into the vector backbones, generating T7-GFPTrap and T7HP-GFPTrap. Both plasmids were sequence-verified using nanopore sequencing.

### IRES Library Construction

Libraries containing candidate viral, cellular, type IV, type VI, and IAPV, Zika, and Dengue variants were synthesized as oligo pools (Twist Bioscience) and 10 ng of each pool was amplified using Q5® High-Fidelity Polymerase (New England Biolabs) in a 25 μl reaction for 12 cycles with the primers IRES-lib-F and IRES-lib-R (Table S8). PCR products and the T7-GFPTrap and T7HP-GFPTrap vectors were digested with HindIII and BglII and ligated to the digested PCR products. Type IV candidate IRES sequences too long to synthesize as oligo pools were synthesized as gene fragments (Twist Bioscience, Table S8), digested with HindIII and BglII, cloned into T7-GFPTrap and T7HP-GFPTrap vectors. The constructs were pooled and added to the Type IV and Type VI IRES libraries cloned from oligo pools. The HCV scanning mutagenesis library was synthesized as two oligo pools (Twist Bioscience), one that contained the mutations in the 5’ end of the sequence (fragment 1) and the other containing the mutations in the 3’ end (fragment 2). Due to sequencing limitations, a five base pair barcode sequence was added just downstream of the HindIII site in fragment 1 oligos. The oligo pools were amplified as previously described with the primers HCV-frag1 F/R and HCV-frag2 F/R (Table S8). The fragment 1 PCR product was digested with HindIII and NheI, the fragment 2 PCR product was digested with NheI and BglII. The wild-type HCV constructs in T7-GFPTrap and T7HP-GFPTrap were digested either with HindIII and NheI or NheI and BglII and the remaining wild-type HCV sequence and vector backbones were gel purified and ligated to the fragment 1 and 2 pools. The ligations were purified using a PCR cleanup column and transformed by electroporation. Transformants were grown overnight in 100 mL LB medium containing ampicillin, and plasmid DNA was extracted for use as a PCR template.

### Construction of GFP-Trap IRES controls

Control constructs (HCV, CrPV, CDK11a, Hoxa9, and Scrambles 1–3) were adapted from previously described vectors^22^. NanoLuc luciferase was removed by digestion with BglII and XbaI and replaced with the GFP-Trap construct (see Construction of GFP-Trap vectors). Three spike-in controls containing randomly scrambled versions of the Eno2 5′UTR (lacking upstream ATG sequences) upstream of GFP-Trap were synthesized as gene fragments (Table S8).

Constructs were pooled into control groups as follows: IRES control group A contained equal amounts of HCV, CrPV, CDK11a, mouse Hoxa9, human Hoxa9, and Scrambles 1–3 in the T7HP-GFPTrap background. IRES control group G contained equal amounts of HCV and CrPV in the T7-GFPTrap background. Spike-in controls were mixed such that spike-in 1 was 100-fold lower than spike-in 3, and spike-in 2 was 10-fold lower than spike-in 3.

### RNA synthesis

To add a poly(A) tail to library pools and individual constructs, 1 ng of plasmid library, control pools, or gene fragments (spike-in controls) was used as a template in a 50 μl PCR reaction with primers phMGFP-T7-F and pHMGFP-3pR. The reverse primer encoded a 70-base poly(A) tail, which was incorporated into the PCR product. A total of 500 ng of PCR product was then used as a template for RNA synthesis using the HiScribe® T7 Quick High Yield RNA Synthesis Kit, following the manufacturer’s instructions. A standard cap analog (m7G(5′)ppp(5′)G) was included in transcription reactions for T7-GFPTrap constructs and spike-in controls, while an A-cap analog (m7G(5′)ppp(5′)A) was used for T7HP-GFPTrap constructs. The resulting RNA was ethanol-precipitated and resuspended in 100 μl of nuclease-free water. RNA integrity was assessed by gel electrophoresis, and concentration was quantified using a Qubit™ RNA High Sensitivity Assay (Invitrogen), according to the manufacturer’s instructions.

### Immunoprecipitation

For each biological replicate, 1×10^6^ HEK293T cells were seeded in 10 cm dishes in 10 mL of DMEM supplemented with 10% FBS. The cells were incubated overnight so that they were 50-70% confluent at the time of transfection. The cells were transfected for 10 hours with 2 μg of A-and G-capped of *in vitro* transcribed RNA pools mixed with 20 μl of Lipofectamine™ MessengerMAX™ (Invitrogen) in 1 mL of Opti-MEM™ (Gibco). Either 8 μl of DMSO (control) or 8 μl of 1.25 mM thapsigargin (ER stress) was added to 500 mL Opti-MEM™ and then added to each plate, for a final concentration of 1 μM thapsigargin for the stress condition. The cells were incubated for 3 hours at 37 °C. The cells were harvested by adding cycloheximide (CHX) to a final concentration of 100 μg/mL and incubating for 10 minutes at 37 °C. The media was removed and 10 mL of ice-cold PBS with 100 μg/mL CHX was added to the dish. The cells were dislodged from the plate using a cell scraper and transferred to a 15 mL conical tube. The cells were centrifuged for 1 minute at 500 × g and the PBS was decanted. The cells were placed on ice and any remaining PBS was removed. The cell pellet was resuspended in 1 mL lysis buffer (50 mM Tris-HCl pH 7.5, 140 mM KCl, 12 mM MgCl_2_, 1% Triton X-100, 1 mM DTT, 100 μg/mL CHX, 1x cOmplete™ EDTA-free protease inhibitor cocktail (Roche), 0.2 U/μl SUPERase-In™ (Invitrogen), and 0.1 U/μl RNasin® Ribonuclease inhibitor (Promega)). The cells were allowed to lyse on ice for 10 minutes and then transferred to 1.5 mL tubes. The lysate was centrifuged at 20,000 × g for 10 minutes at 4 °C and the supernatant was removed to a new tube. A 50 μl sample of lysate was saved in 500 μl of TRIzol™(Invitrogen) for later RNA extraction. A spike-in only transfected lysate (unstressed) was prepared simultaneously and added to each lysate to 1% of the total volume. Anti-HA magnetic beads (Pierce) were prepared by placing 20 μl of beads per sample into 1.5 mL tubes. The storage buffer was removed, and the beads were washed with 200 μl of lysis buffer for 10 minutes. Each lysate was divided into replicate IP samples, so that 450 μl of lysate was added to 20 μl of magnetic beads and incubated at 4 °C for 2 hours. The beads were washed 3 times with 800 μl of a high salt buffer (50 mM Tris-HCl pH 7.5, 300 mM KCl, 12 mM MgCl_2_, 1% Triton X-100, 0.5 mM DTT, 100 μg/mL CHX, 1× complete™ EDTA-free protease inhibitor cocktail (Roche), 0.2 U/μl SUPERase-In™ (Invitrogen), and 0.1 U/μl RNasin® Ribonuclease inhibitor (Promega)) for 10 minutes per wash at 37 °C. The beads were then washed twice for 10 minutes with 200 μl of lysis buffer at 37 °C. The beads were resuspended in 250 μl of lysis buffer and 25 U of AcTEV™ protease (Invitrogen) were added and the beads were incubated on a tube rotator for 16 hours at 4 °C. The samples were placed on a magnet and the supernatant was removed and placed into a 1.5 mL tube containing 750 μl of TRIzol™, and RNA was extracted following the manufacturer’s instructions. Input RNA (IN RNA) was precipitated and resuspended in 50 μl of nuclease-free water while RNA from the immunoprecipitation (IP RNA) was resuspended in 23 μl.

### Sequencing library preparation

The reverse transcription (RT) primers, IP-BC 9-11 (Table S8) contained either 9, 10, or 11 random bases so that a unique molecular index (UMI) could be incorporated into each cDNA synthesized from the input and IP RNA. In a 20 μl reaction using SuperScript™ (Invitrogen) reverse transcriptase, 100 ng of input RNA and 11 μl of IP RNA was reverse transcribed at 55°C for 30 minutes. The cDNA was purified using a 1X concentration of magnetic PCR purification beads according to the manufacturer’s instructions and the cDNA was resuspended in 10 μl of nuclease free water. The cDNA was amplified for 6 cycles using the primers IPN0-N6F and IP-RT-R (Table S8) in a 25 μl reaction using Q5® High-Fidelity DNA Polymerase (New England Biolabs). The PCR product was purified using 1X concentration of magnetic PCR purification beads and eluted in 16 μl. To add on Illumina barcodes and sequences, 2 μl of the first PCR reaction was amplified for 15 cycles in a 25 μl reaction using Q5® Polymerase with dual indexed primers (NEBNext® Multiplex Oligos). The PCR product was purified with 1X concentration of magnetic PCR purification beads and eluted in 15 μl nuclease-free water. The libraries were sequenced 2 × 150 bp reads on an Illumina sequencer.

### Sequence data processing and analysis

Raw sequence data were processed to quantify unique (UMI) reporter construct reads from input and immunoprecipitated samples as follows. Read pairs were first merged using flash (version 1.2.11; -z -t 3 -O -M 150). This resulted in single reads for short IRES candidates and unmerged read pairs for larger candidates. Reads were then processed to collapse UMI duplicates using two perl scripts, <u>umi-collapse-singles.pl</u> and <u>umi-collapse-pairs.pl</u>) for merged single reads and unmerged read pairs, respectively. These scripts removed sequences corresponding to the TRAP reporter protein and appended the UMI sequence for each read or readpair to the readIDs. Notably, this only partially collapses UMIs, as sequence duplicates with different illumina library stagger lengths remain present at this stage. Reads were then split into “a-cap” (IRES) or “g-cap” (CDI) groups using cutadapt (version 4.5). For single reads, options - j8, -n 2 --trimmed-only -e 0 were used. The common adapter “AGATCTGTGATTAAG” was first removed, and the resulting file was trimmed to identify a-cap reads that carried the “<u>AGTCCGCATCC</u>AAGCTT” adapter and g-cap reads carrying the “<u>TCCAAGCGATC</u>AAGCTT” adapter. For read pairs, options --pair-adapters --pair-filter both -j 8 --trimmed-only -e 0.07 were used. A-cap reads were identified by trimming with adapters -g CTTAATCACAGATCT -G <u>AGTCCGCATCC</u>AAGCTT, while g-cap reads were identified by trimming adapters -g CTTAATCACAGATCT -G <u>TCCAAGCGATC</u>AAGCTT. Merged single and unmerged paired reads were aligned to the a target database of IRES constructs using hisat2 (version 2.1.0, options --no-spliced-alignment). To accommodate larger insert sizes in Type IV IRESes, the -X 550 option was included. The number of reads aligning to each IRES construct was tabulated using a custom perl script (IP_bam_counter_both.pl), which resolved remaining UMI duplicates by requiring unique UMIs for all reads that align to the same IRES construct. The resulting a-cap and g-cap readcounts for each IRES construct in input and IP samples were compiled into tables for downstream analyses.

The Pearson correlations among all technical replicates were high (>0.99 Figure S1); therefore, read counts were combined by summing across replicates for downstream analysis. Constructs with fewer than 10 total reads in any input library were removed from further analysis.

For Pearson correlation analysis among biological replicates, constructs with zero reads in any IP library replicate were excluded. Reads per million (RPM) were then calculated, and translational efficiency (TE) was defined as the ratio of RPM in the IP library to RPM in the input library. Pearson correlations among biological replicates were also high (>0.94); therefore, read counts were combined by summing across biological replicates. For IP libraries, zero counts were replaced with a small pseudocount (0.000001) prior to RPM calculation. TE was subsequently calculated as IP/Input.

### mRNA Luciferase assays

Test sequences were synthesized as gene fragments (Twist Bioscience, Table S8). Gene fragments containing one construct were digested with HindIII and BglII and cloned into T7-Nluc and T7HP-Nluc^22^. In cases where the constructs were too short to synthesize as individual constructs, constructs were combined as one gene fragment and then PCR amplified, digested with HindIII and BglII, and cloned into the luciferase vectors. All constructs were sequence verified by nanopore sequencing. The luciferase reporter constructs were amplified with the primers phMGFP-T7-F and phMGFP-GFP3p-R and the resulting PCR products were used as templates for T7 transcription. 250 ng of the PCR product was used as a template for RNA synthesis, using the HiScribe® T7 Quick High Yield RNA Synthesis kit (New England Biolabs) according to the manufacturer’s instructions. A standard cap analog, m^7^G(5’)ppp(5’)G, was incorporated into the transcription reaction for T7-nluc constructs, and an A-cap analog, m^7^G(5’)ppp(5’)A, was incorporated into the transcription reaction for T7HP-nluc constructs. The resulting RNA was ethanol precipitated and resuspended in 100 μl of nuclease-free water. The RNA integrity was verified by gel electrophoresis and quantified using a Qubit™ RNA high sensitivity assay (Invitrogen) according to the manufacturer’s instructions. HEK293T cells were plated at 1 × 10^4^ per well in a 96 well plate, in 100 μl of DMEM supplemented with 10% FBS. The cells were incubated overnight at 37°C and transfected at 50-70% confluency. For S2 and Aag2 cells transfections, 1 × 10^5^ and 1 × 10^4^ cells in 100 μl of Schneider’s Drosophila Medium supplemented with 10% FBS were added to each well of a 96 well plate. For each well, 30 ng of each UTR-N-luc RNA was co-transfected with 10 ng of Firefly luciferase (F-luc) control RNA using Lipofectamine™ MessengerMAX™ Transfection Reagent (Invitrogen) for four hours. Six technical replicates were performed for each construct. The chemical stressors (20 nM rapamycin and 50 μM 4EGI-1) were added to the wells 1 hour after the transfection with DMSO as a negative control. Prior to mRNA transfections as described above, for the viral 2A protease stress, 100 ng of viral protease 2A vector was transfected per well using Lipofectamine™ 3000 for 44 hours. The Nano-Glo® Dual-Luciferase® Reporter Assay System (Promega) was used to measure N-luc and F-luc luminescence, according to the manufacturer’s instructions, on a Tecan microplate reader (Table S9).

### Tornado-split-nLuc vector luciferase assays

Candidate IRES sequences were synthesized as gene fragments (Twist Bioscience, Table S8) and amplified with primers that added EcoRI and BsiWI sites (Table S8). The PCR products and pcDNA3.1+-Tornado-split-nLuc vector^38^ (Addgene #212612) were digested with EcoRI and BsiWI, and ligated. All constructs were sequence-verified by nanopore sequencing. Positive (EMCV, HCV, CrPV) and negative (scramble 1) controls were previously described^22^. HEK293T cells were seeded at a density of 2.0 x 10^3^ cells per well of a 96 well plate in 100 μl of DMEM / 10% FBS and incubated overnight at 37°C. HEK293T cells were transfected for 24 hours with 75 ng of each reporter construct per well with 45 ng of a control vector expressing Firefly luciferase from the mouse *Hoxa5* promoter^20^ using LipofectamineTM 3000 transfection reagent (Invitrogen). Six replicates were performed for each construct. The Nano-Glo® Dual-Luciferase® Reporter Assay System (Promega) was used to measure N-luc and F-luc luminescence, according to the manufacturer’s instructions, on a Tecan microplate reader. Data were processed by subtraction of luminescence values from untransfected negative control wells and N-luc / F-luc ratios were calculated for each replicate.

### Viral protease 2A construct cloning

The EMCV IRES sequences were amplified from the EMCV-T7-nluc construct using the primers EMCV-Ext-SacI-F and Vector-BglII-R (Table S8). The PCR products were digested with SacI and BglII and inserted in the T7-nluc vector to replace the UTR sequence. Human rhinovirus A-16 protease 2A (HRV2A^pro^) wild-type and an inactive mutant sequence (HRV2A^pro^ Cys106Ala^59^) were synthesized as gene fragments (Table S8) with GFP upstream of VP16 followed by the 2A protease sequence. The gene fragments and previously described EMCV-nluc vector were digested with BglII and XbaI and n-luc was replaced with the viral 2A protease sequences, resulting in the constructs EMCV-HRV2A^pro^ and EMCV-HRV2A^pro^ C106A.

### Polysome profiling

To assess the translational response to various stressors, HEK293T cells were stressed by 1 μM thapsigargin, 20 nM rapamycin, 50 μM 4EGI1, and by expression of viral protease 2A in the cells. For all the stressors, 1 × 10^6^ cells were seeded in 10 cm dishes in 10 mL of DMEM supplemented with 10% FBS. The cells were incubated overnight at 37°C. The chemical stressors were added for 3 hours using DMSO only as a no stress control. For the viral protease 2A stress, the cells were transfected with 10 μg of either EMCV-HRV2A^pro^ or EMCV-HRV2A^pro^ U106A plasmid DNA for 48 hours using Lipofectamine™ 3000 transfection reagent. To harvest the cells, the media was removed, and the cells were resuspended in 10 mL of ice-cold PBS supplemented with 100 μg/mL cycloheximide by using a cell scraper to dislodge the cells. The cell suspensions were transferred to 15 mL conical tubes and pelleted for 1 minute at 500 x g. The supernatant was decanted, and the cells were resuspended in 500 μl of PLB (50 mM Tris-HCl pH 7.5, 140 mM KCl, 12 mM MgCl_2_, 1% Triton X-100, 1 mM DTT, cOmplete™ EDTA-free protease inhibitor cocktail (Roche), 100 mg/mL cycloheximide, 0.2 U/μl SUPERase-In™ (Invitrogen), and 0.1 U/μl RNasin® Ribonuclease inhibitor (Promega)). The cells were allowed to lyse on ice for 10 minutes and then transferred to 1.5 mL tubes. The lysate was centrifuged at 20,000 x g for 10 minutes at 4°C and the supernatant was removed to a new tube. The lysates were centrifuged over a 10–50% sucrose gradient (in PLB without Triton X-100) and the polysome profiles were recorded across the gradient using a Biocomp polysome fractionator.

## Data availability

All primary sequencing data have been deposited at the Sequence Read Archive data repository (https://www.ncbi.nlm.nih.gov/sra/) under accession number PRJNA1453056.

## Acknowledgements

The authors thank members of the McManus lab for helpful comments and suggestions. We also thank Shawn Lyons for providing the Aag2 mosquito cell line and advice on its usage, and Deepika Vasudevan for providing S2 cells. This work was supported by NIH grant R35GM145317 to CJM.

---

**Figure S1.**
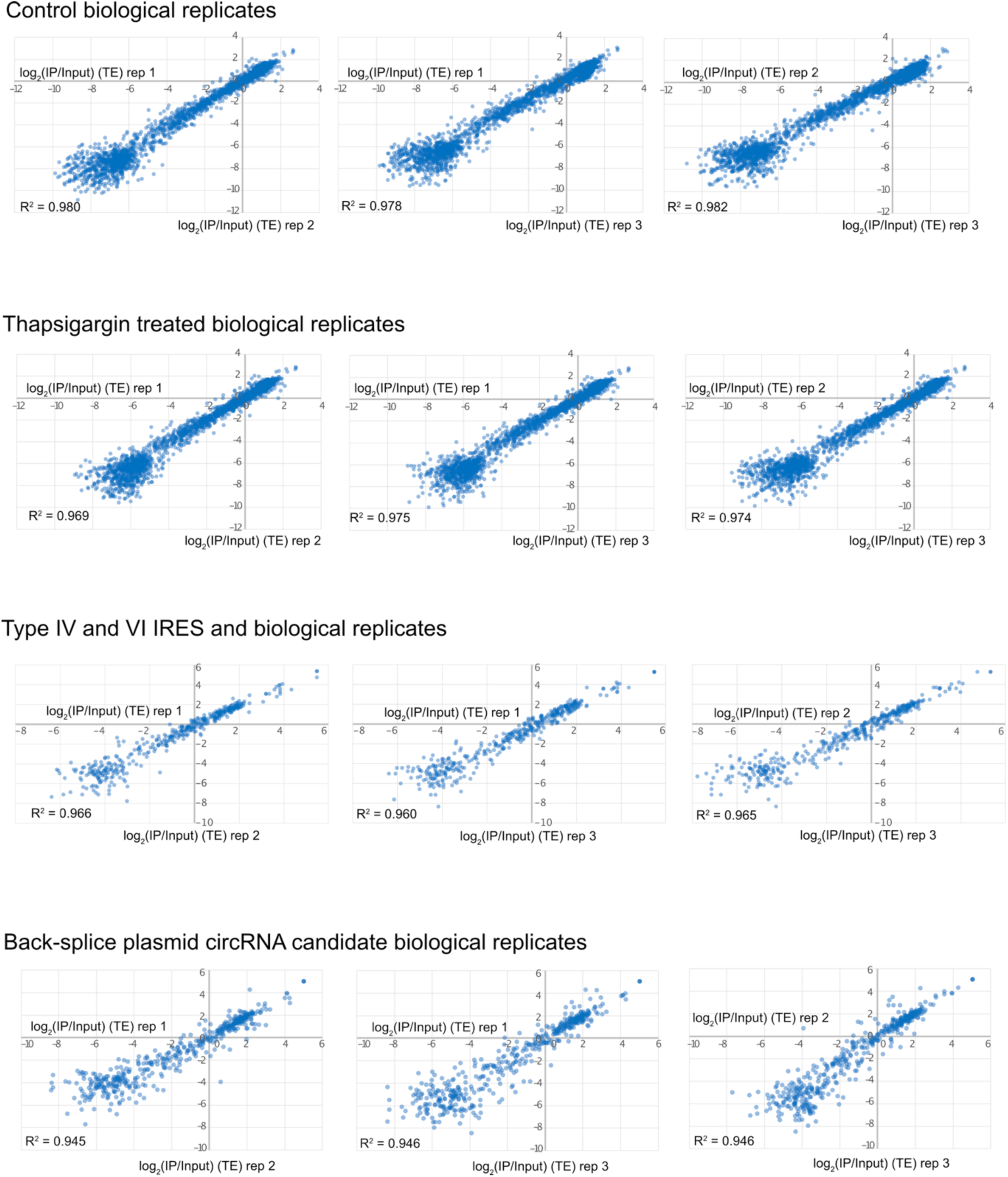
Comparison of biological IRES-TrAPPr replicates. Scatter plots show combined TE estimates for CDI and IRES activity for the experiments described in this manuscript. In all cases, there was strong correlation between replicates (R^2^ >= 0.95).

**Figure S2.**
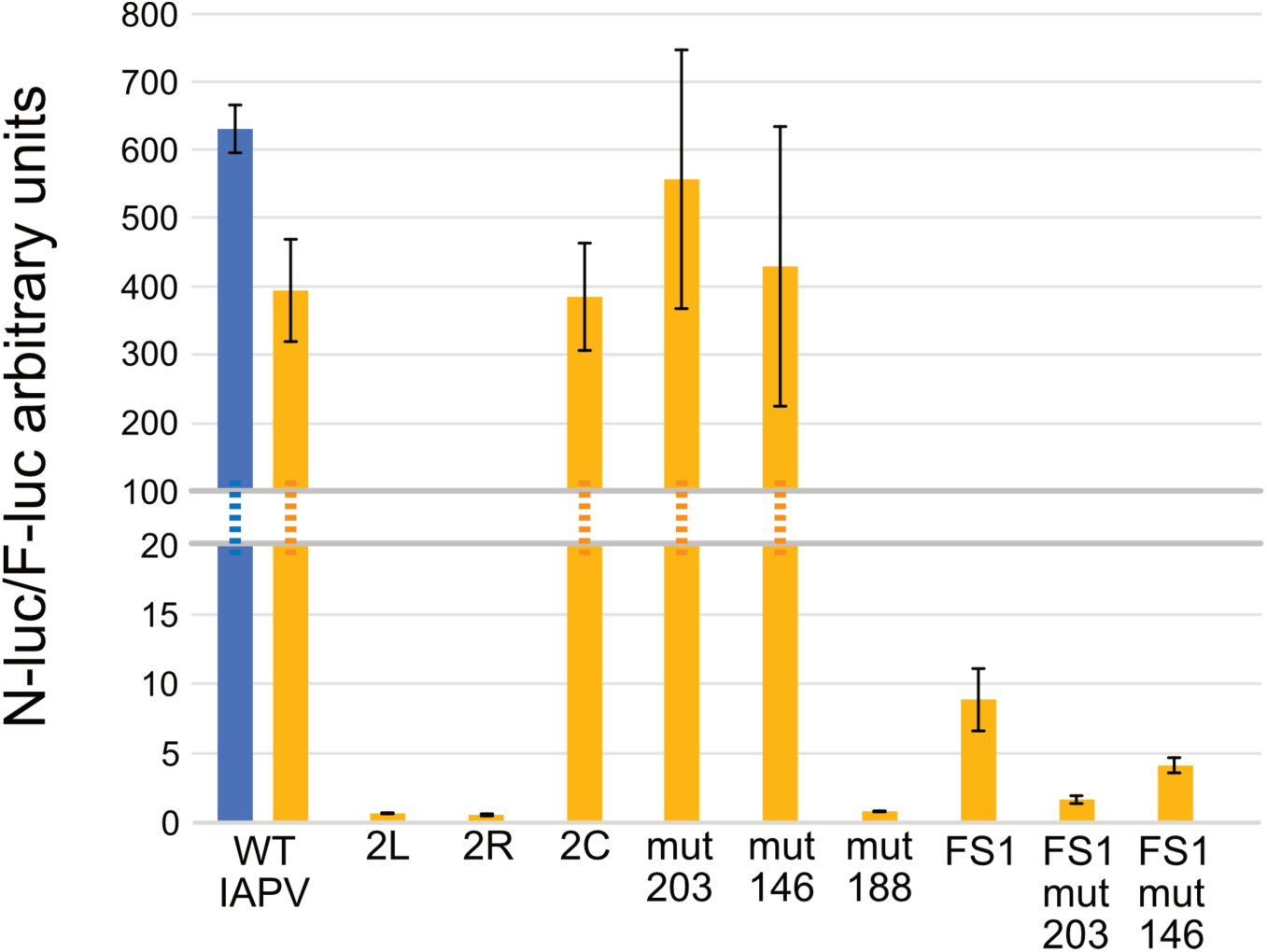
Luciferase reporter validation of IAPV variants. G-cap and A-cap (IRES) n-Luc reporter RNAs were transfected into HEK293T cells along with a common F-luc control (see methods). Mutations that disrupted basepairing in the second pairing region (2L and 2R) nearly eliminated IRES activity, while their combination (2C) restores basepairing and IRES activity. Mutations at positions 203 and 146 increased IRES activity in IRES-TrAPPr and individual reporters, however this was not statistically significant in individual assays. Mutation 188 decreased IRES activity in both IRES-TrAPPr and validation reporters. The FS1 mutation represents IRES activity in the +1 reading frame (n-Luc cloned in +1). FS1 mut 203 and mut 146 are +1 reading frame reporters with mutations at 203 and 146, both of which decrease +1 translation.

**Figure S3.**
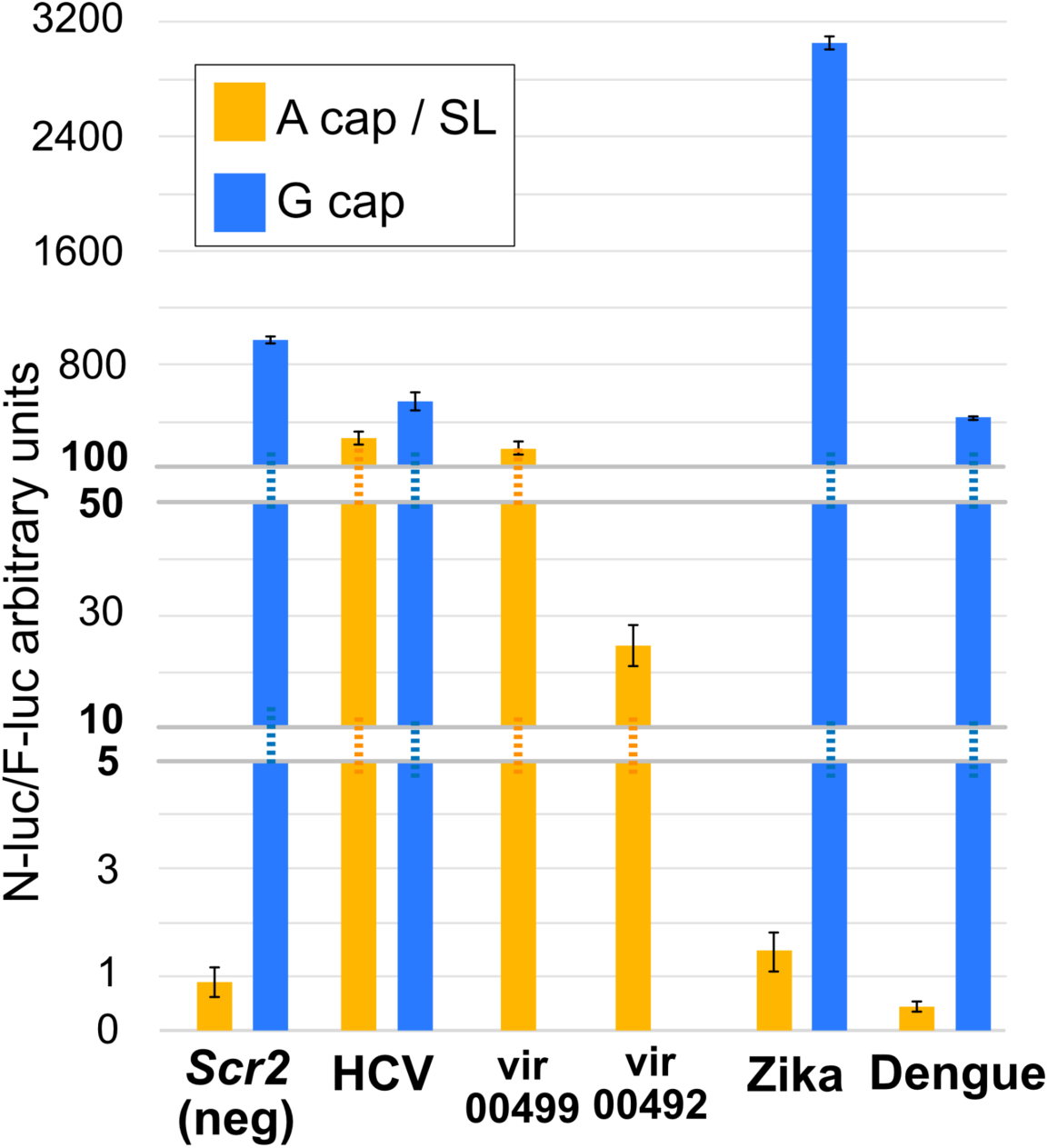
Luciferase reporter validation of viral IRES candidates. G-cap and A-cap (IRES) n-Luc reporter RNAs were transfected into HEK293T cells along with a common F-luc control (see methods). The two positive IRESbase candidates had strong IRES activity, while the 5’ UTRs of Zika and Dengue virus had negligible activity.

**Figure S4.**
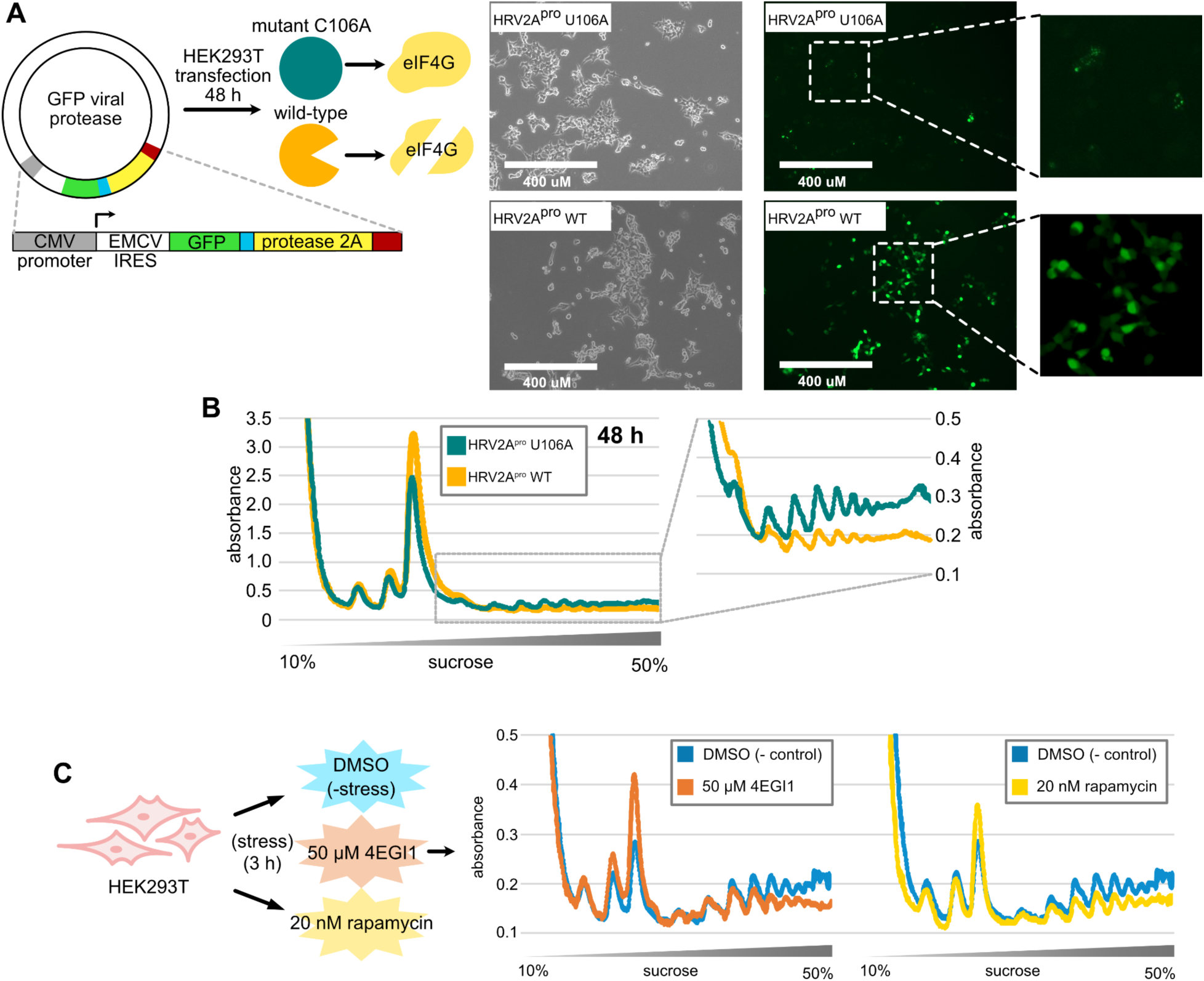
Expression of 2A protease and treatment with translational inhibitors induces translational stress. **A.** Plasmids encoding wildtype and nonfunctional U106A (Cys106Ala) Human Rhinovirus protease 2A fused to GFP were transfected into HEK293T cells. The blue segment in the diagram between GFP and the 2A protease is a 28 amino acid peptide recognized by the 2A protease for auto excision from GFP. Microscopy images of transfected cells show stable expression of GFP fused to wildtype, but not 2A mutant protease. **B.** Polysome gradient fractionation shows expression of 2A protease reduces polysomes and accumulates monosomes. **C.** Cells were separately treated with translation inhibitors 4EGI1 and rapamycin (left), which also reduced polysome accumulation and increased monosome abundance (right).

